# Processivity of the monomeric KLP-6 kinesin and a Brownian ratchet model with symmetric potentials

**DOI:** 10.1101/2024.08.06.606760

**Authors:** Tomoki Kita, Kazuo Sasaki, Shinsuke Niwa

## Abstract

Most kinesin molecular motors dimerize to move processively and efficiently along microtubules; however, some can maintain processivity even in a monomeric state. Previous studies have suggested that asymmetric potentials between the motor domain and microtubules underlie this motility. In this study, we demonstrate that the kinesin-3 family motor protein KLP-6 can move along microtubules as a monomer upon release of autoinhibition. This motility can be explained by a change in length between the head and tail, rather than by asymmetric potentials. Using mass photometry and single-molecule assays, we confirmed that activated full-length KLP-6 is monomeric both in solution and on microtubules. KLP-6 possesses a microtubule-binding tail domain, and its motor domain does not exhibit biased movement, indicating that the tail domain is crucial for the processive movement of monomeric KLP-6. We developed a mathematical model to explain the unidirectional movement of monomeric KLP-6. Our model concludes that a slight conformational change driven by neck-linker docking in the motor domain enables the monomeric kinesin to move unidirectionally if a second microtubule-binding domain exists.

**SIGNIFICANCE:** Kinesin molecular motors are designed to move efficiently using two heads. Studying these biological molecular motors provides valuable insights into the mechanisms that generate unidirectional movements amidst intense thermal fluctuations. This study reveals that the monomeric kinesin-3 KLP-6 can move along microtubules through interactions with its tail domain. The proposed Brownian ratchet model explains this movement by considering a change in stalk length caused by neck-linker docking rather than asymmetric potentials. This model suggests that a slight conformational change can achieve robust processive movement of kinesin. These findings have significant implications for understanding Brownian ratchet motors and designing rational artificial molecular motors.

## INTRODUCTION

Typical kinesins are homodimers composed of two motor domains that move along microtubules in a manner similar to how we use our legs when we walk. They convert chemical energy from ATP hydrolysis into mechanical work, facilitating cargo transport (1). Conventional dimeric kinesin, known as kinesin-1, is remarkably efficient, taking about 100 forward steps per second and rarely stepping backward under no load (2–5). The precise walking mechanism of dimeric kinesin has garnered significant attention as a model for understanding robust molecular motor dynamics amidst intense thermal fluctuations (6).

Some kinesins have been reported to exhibit processivity along microtubules even in a monomeric state. KIF1A, a member of the kinesin-3 family motor proteins, transports synaptic vesicle precursors in axons (7–9). Initially considered a unique monomeric motor (7, 10), KIF1A has subsequently been shown to form homodimers when activated (11, 12). KIF1A possesses a lysine-rich insertion in the L12 loop, known as the “K-loop,” which allows its truncated monomeric head to undergo biased Brownian motion on microtubules even in a monomeric state (10). The Brownian ratchet model effectively explains this monomeric movement (10, 13–16). A later study demonstrated that truncated monomeric kinesin-1 (K351) can also move processively along microtubules under low-ionic conditions (17). Surprisingly, attaching a non-catalytic microtubule-binding protein, BDTC (a 1.3-S subunit of *Propionibacterium shermanii* transcarboxylase), to the C-terminal region of K351 accelerated the motor speed by four times and reduced the diffusion coefficient to one-third (17). Similarly, budding yeast kinesin-14 Kar3 heterodimerizes with a non-catalytic motor domain, either Cik1 or Vik1 (18), and moves processively on microtubules, even though monomeric Kar3 is not processive (19). Other studies have revealed that a non-processive motor head can become processive through dimerization with a processive partner head (20–22). The mechanism that allows a non-processive motor to transition to a processive state through the attachment of another microtubule-binding domain remains elusive, as the prevailing Brownian ratchet model or models for dimeric kinesin cannot adequately explain their motilities (10, 13–16, 23, 24).

Interestingly, while some kinesin-3 family members form both monomers and dimers in solution due to their autoinhibition (12, 25), the invertebrate-specific kinesin-3 KLP-6 forms only monomers (12, 26). KLP-6 transports a mechanosensory receptor complex in male cilia (27, 28). We and another group have shown that full-length KLP-6 is a strongly autoinhibited monomeric protein that does not move processively along microtubules (12, 26). Its MBS and MATH tail domains can bind to microtubules, allowing autoinhibited full-length KLP-6 to remain attached to them (12). Deleting the MBS and MATH domains (KLP-6(1-587)) and releasing autoinhibition (KLP-6(1-587)(D458A)) did not facilitate the dimerization or processive movement along microtubules (12). Although the truncated motor domain remained monomeric and non-processive, artificial dimerization using a leucine zipper domain enabled the motor domains to achieve processivity (12). However, the dimerization and processive movement of full-length KLP-6 have not yet been confirmed.

In this study, we show that the activated form of full-length KLP-6 moves processively along microtubules even in a monomeric state. We propose a Brownian ratchet model to explain how interactions between the tail domain and microtubules confer processivity to the non-processive motor domain. Our model suggests that neck-linker docking in the motor domain is essential for this movement, indicating a mechanism by which kinesin motor domains acquire processivity when combined with a second microtubule-binding domain.

## MATERIALS AND METHODS

### Plasmid

The plasmid KLP-6(D458A)::sfGFP::Strep-tag II was generated by PCR-based mutagenesis using KOD plus neo DNA polymerase (TOYOBO), based on the KLP-6::sfGFP::Strep-tag II vector (12). To create KLP-6(D458A)::mScarlet::Strep-tag II, the plasmids KLP-6(D458A)::sfGFP::Strep-tag II and Xkid::mScarlet::Strep-tag II (29) were digested with HindIII/NotI and ligated using Ligation High Ver.2 (TOYOBO).

### Purification of recombinant proteins

KLP-6::sfGFP::Strep-tag II, KLP-6(1-390)::sfGFP::Strep-tag II, and KLP-6(580-928)::sfGFP::Strep-tag II were previously purified (12) and reanalyzed in this study. We purified KLP-6(D458A)::sfGFP::Strep-tag II and KLP-6(D458A)::mScarlet::Streptag II as described (12) (see Section S1 of the supporting materials and methods for more detail). The proteins were expressed in Sf9 cells (Thermo Fisher Scientific) infected with P1 baculovirus. Affinity purification was performed using Streptactin-XT resin (IBA Lifesciences). Eluted fractions were further separated using an NGC chromatography system (Bio-Rad) equipped with a Superdex 200 Increase 10/300 GL column (Cytiva).

### Mass photometry

Proteins obtained from the peak fractions in the SEC analysis were pooled, snap-frozen, and stored until measurement. Prior to measurement, the proteins were thawed and diluted to a final concentration 20 nM in dilution buffer (50 mM HEPES-KOH, pH 7.5, 150 mM KCH3COO, 2 mM MgSO4, 1 mM EGTA). Mass photometry was performed using a Refeyn OneMP mass photometer (Refeyn Ltd, Oxford, UK) and Refeyn AcquireMP version 2.3 software, with default parameters set by Refeyn AcquireMP. Bovine serum albumin (BSA) was used as a control to determine the molecular weight. The results were subsequently analyzed using Refeyn DiscoverMP version 2.3, and graphs were prepared to visualize the data.

### Total internal reflection fluorescence single-molecule motility assays

Total internal reflection fluorescence (TIRF) assays were performed as described (12) (see Section S2 of the supporting materials and methods for more detail). All assays were performed in assay buffer (90 mM HEPES-KOH pH 7.4, 50 mM KCH_3_COO, 2 mM Mg(CH_3_COO)_2_, 1 mM EGTA, 10% glycerol, 0.1 mg/mL biotin-BSA, 0.2 mg/mL kappa-casein, 0.5% Pluronic F127, nucleotide of interest (ATP, ADP, or AMP-PNP) diluted to indicated concentrations, and an oxygen scavenging system composed of PCA/PCD/Trolox). Purified motor proteins were diluted to concentrations suitable for observing single particles (2-150 pM) in the assay buffer. Diluted KIF5A(1-416)-mScarlet was added to the assay buffer to determine the polarity of the microtubules. The solution was then introduced into the glass chamber. An ECLIPSE Ti2-E microscope equipped with a CFI Apochromat TIRF 100XC Oil objective lens (1.49 NA), an Andor iXon Life 897 camera, and a Ti2-LAPP illumination system (Nikon, Tokyo, Japan) was used to observe single-molecule motility. NIS-Elements AR software version 5.2 (Nikon) was used to control the system. KLP-6 proteins in the presence of 2 mM MgATP, 2 mM MgAMP-PNP, or 5 mM MgADP treated with hexokinase (30) were recorded at a frame rate of 10 frames per second (fps). For fluorescence intensity analysis, to reduce intense motor fluctuations and obtain clear fluorescence, the moving particles were observed in the presence of 2 *μ*M MgATP at a frame rate of 5 fps. In dual-color single-molecule assays, sequential images of the 488 and 561 channels were taken at a frame rate of 0.7 fps.

### Data analysis and graph preparation

All analyses were conducted using kymographs generated by ImageJ software. Motors that remained on microtubules for over 1.0 s (10 pixels) were analyzed. The instantaneous velocity was calculated from the pairwise distances for a window size of 1.0 s (17, 31). The mean square displacement (MSD) *ρ*(*τ*) was plotted by averaging squared displacements for intervals *τ* (32). The diffusion constant and the drift velocity were determined by fitting *ρ*(*τ*) with *ρ*(*τ*) = 2*Dτ* + *v*^2^*τ*^2^ + *C* or *ρ*(*τ*) = 2*Dτ* + *C* (*D*, diffusion coefficient; *v*, drift velocity; *C*, constant) to the first several time intervals of the obtained MSD plots. For KLP-6-GFP in the AMP-PNP solution, periods with displacements of 5 pixels or more were analyzed as the weak microtubule binding state, while other periods were analyzed as the strong microtubule binding state. For fluorescence intensity analysis, the background-subtracted maximum fluorescence intensity during motor binding to microtubules was obtained. Statistical analyses were performed using GraphPad Prism version 9. Graphs were prepared using Python, exported in PNG format, and aligned using Adobe Illustrator 2021.

## RESULTS

### Releasing autoinhibition enables full-length KLP-6 to become processive on microtubules

We quantified the motility of KLP-6 proteins on microtubules by measuring instantaneous velocity and MSD using TIRF microscopy. All proteins were fused with sfGFP at their C-terminus, and full-length KLP-6 (KLP-6-GFP), its motor domain (KLP-6(1-390)-GFP), and its MBS-MATH domains (KLP-6(580–928)-GFP) were examined (Fig. 1A). KLP-6-GFP and KLP-6(1-390)-GFP did not show biased movements (the *p* values for the hypothesis that the mean equals zero were 0.4272 and 0.5881 for KLP-6-GFP and KLP-6(1-390)-GFP, respectively; Student’s t-test) (Fig. 1B and C). The diffusion coefficient *D*_ATP_ of KLP-6-GFP is closer to that of KLP-6(580-928)-GFP, which is the largest of all analyzed proteins, rather than that of KLP-6(1-390)-GFP (Figs. 1D and Table 1). This result suggests that KLP-6-GFP spends most of its time in an autoinhibited state on microtubules, and its motor domain rarely binds to them.

**Table 1:** Summary of single-molecule motility assays for KLP-6 proteins.

| Construct | Oligomeric state | Instantaneous $v$ (nm s<sup>-1</sup>) | Drift $v$ (nm s<sup>-1</sup>) | $D_{\text{ATP}}$ ($\mu\text{m}^2 \text{s}^{-1}$) | $D_{\text{ADP}}$ ($\mu\text{m}^2 \text{s}^{-1}$) | $D_{\text{AMP-PNP}}$ ($\mu\text{m}^2 \text{s}^{-1}$) |
| --- | --- | --- | --- | --- | --- | --- |
| KLP-6-GFP | Monomer | $-8.9 \pm 11.2$ (0.4272) | 0 | $0.146 \pm 0.004$ | $0.081 \pm 0.005$ | 0(s) / $0.127 \pm 0.022$(w) |
| KLP-6(1-390)-GFP | Monomer | $5.5 \pm 10.2$ (0.5881) | 0 | $0.034 \pm 0.002$ | $0.038 \pm 0.008$ | 0 |
| KLP-6(D458A)-GFP | Monomer | $67.2 \pm 4.8$ (<0.0001) | $69.4 \pm 24.1$ | $0.009 \pm 0.004$ | $0.025 \pm 0.001$ | 0 |
| KLP-6(580-928)-GFP | Monomer | nd | 0 | $0.189 \pm 0.010$ | nd | nd |

**Figure 1:**
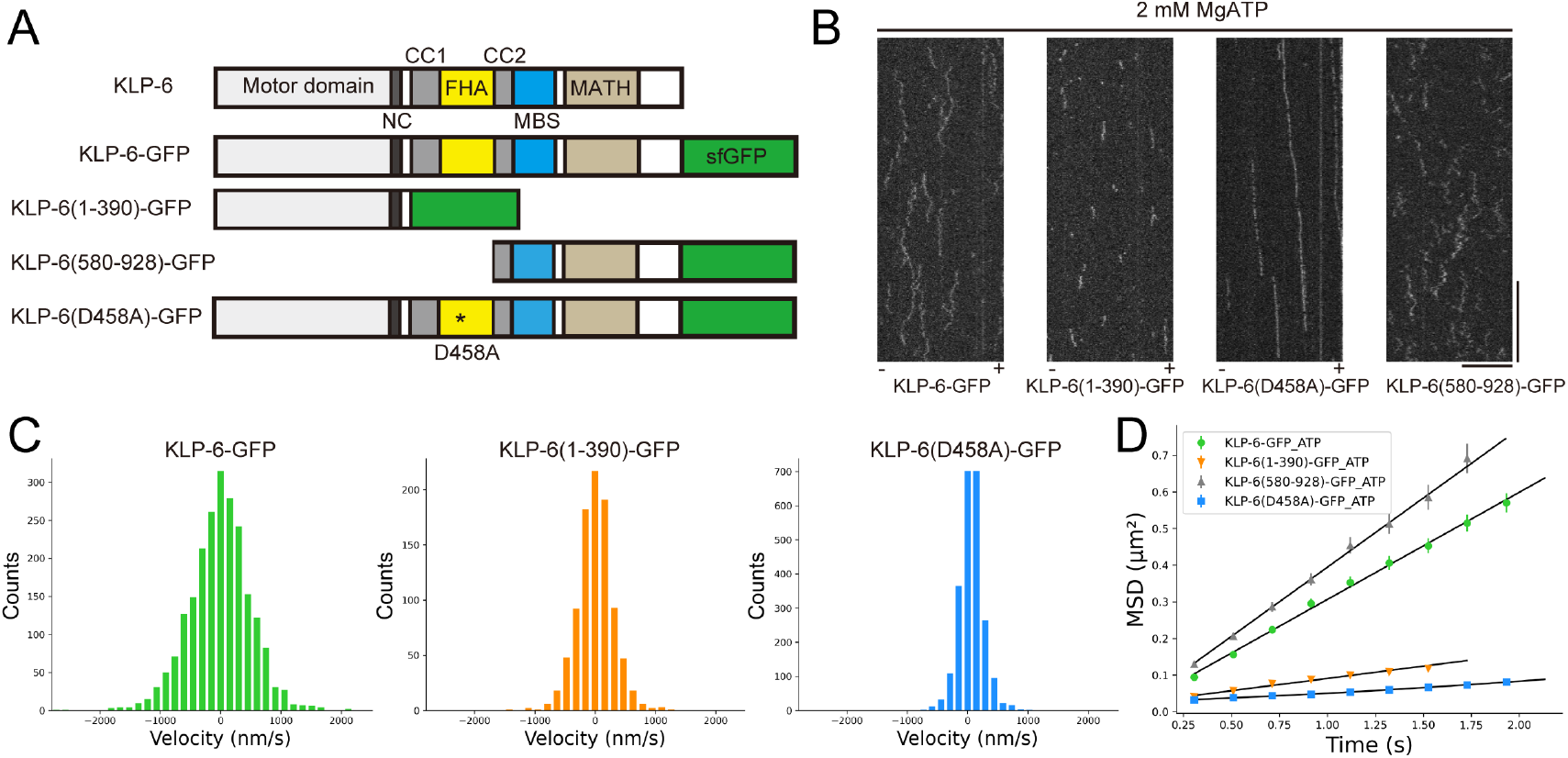
Movements of KLP-6 proteins on microtubules in the presence of ATP. (A) Schematic drawing of the domain organization of KLP-6 analyzed. All proteins are fused with sfGFP. NC, neck coiled-coil domain; CC1, coiled-coil 1 domain; FHA, forkhead-associated domain; CC2, coiled-coil 2 domain; MBS, membrane-associated guanylate kinase homolog (MAGUK)-binding stalk domain; MATH, meprin and TRAF homology domain. D458A is the mutation located in the FHA domain. (B) Representative kymographs showing the motility of the KLP-6 proteins in the presence of 2 mM MgATP. Plus (+) and minus (-) symbols indicate the polarity of microtubules. Horizontal and vertical bars represent 10 *μ*m and 10 s, respectively. (C) Histograms of instantaneous velocities of KLP-6-GFP, KLP-6(1-390), and KLP-6(D458A)-GFP. (D) MSD plots of KLP-6-GFP, KLP-6(1-390), KLP-6(580-928), and KLP-6(D458A)-GFP in the ATP solution with fitted quadratic curves. Each plot represents mean ± SEM. 280-508 particles were analyzed.

To quantify the motility of activated KLP-6 on microtubules, we next analyzed KLP-6(D458A)-GFP, a mutant with disrupted autoinhibition between the FHA domain and motor domain (26) (Fig. 1A). As a result, *D*_ATP_ of KLP-6(D458A)-GFP was significantly reduced compared to KLP-6-GFP, and KLP-6(D458A)-GFP showed a clear bias toward the plus end of microtubules (*p* < 0.0001) (Figs. 1B-D, Table 1, and Movie S1). In our previous study, KLP-6(1-587)(D458A) showed no biased movements (12), indicating that the microtubule-binding tail domains (MBS and MATH domains) are essential for the processivity of monomeric KLP-6(D458A).

### The tail domain suppresses the diffusion of activated KLP-6 in a weak microtubule binding state

It is known that typical kinesins bind strongly to microtubules in the ATP state and weakly in the ADP state (23). Here, we examined the nucleotide-dependent microtubule affinity of KLP-6 and the contribution of its tail domain. In the presence of the non-hydrolyzable ATP analog, AMP-PNP, we observed strong binding of KLP-6-GFP, KLP-6(1-390)-GFP, and KLP-6(D458A)-GFP to microtubules (Fig. 2A and C). Interestingly, some KLP-6-GFP molecules exhibited highly diffusive movements similar to those observed in the presence of ATP before they either strongly bound to or detached from microtubules (Fig. 2A, C, and Table 1). Since ADP is necessary for the autoinhibition of full-length KLP-6 (26), our result suggests that some KLP-6-GFP molecules still contain ADP and remain autoinhibited even in the presence of excess AMP-PNP.

**Figure 2:**
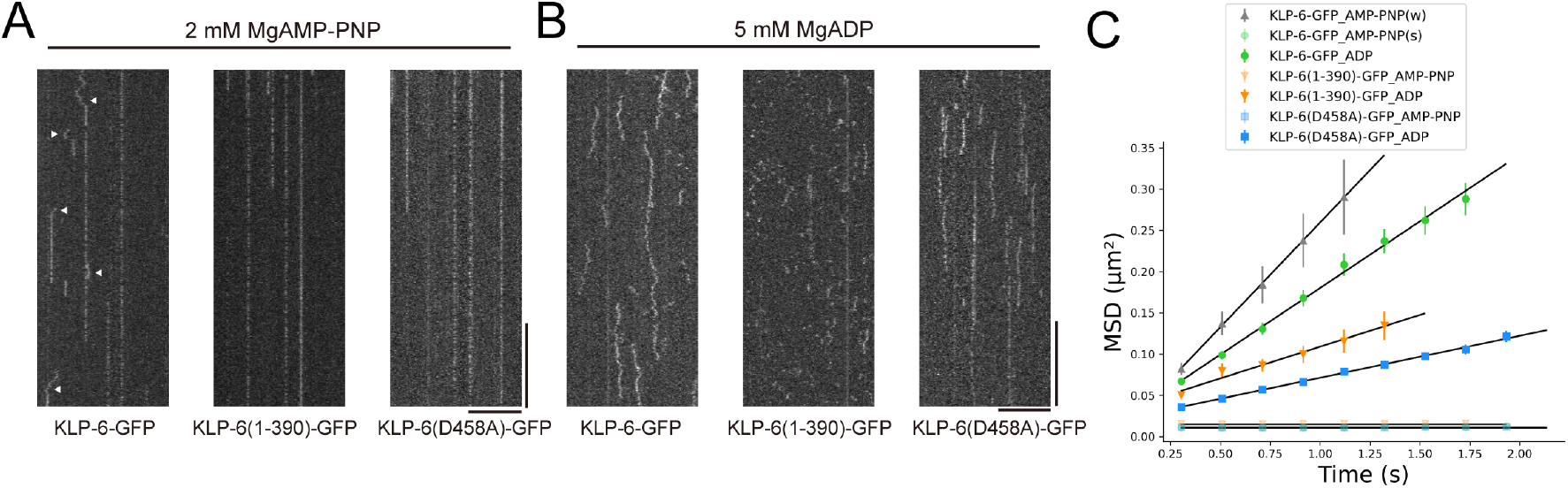
Movements of KLP-6 proteins on microtubules in the presence of AMP-PNP or ADP. (A) Representative kymographs showing the motility of KLP-6 proteins in the presence of 2 mM MgAMP-PNP. Arrowheads indicate examples of the highly diffusive movements of KLP-6-GFP before strong binding to or detachment from microtubules. Horizontal and vertical bars represent 10 *μ*m and 10 s, respectively. (B) Representative kymographs showing the motility of KLP-6 proteins in the presence of 5 mM MgADP. Horizontal and vertical bars represent 10 *μ*m and 10 s, respectively. (C) MSD plots of KLP-6-GFP, KLP-6(1-390)-GFP, and KLP-6(D458A)-GFP in the AMP-PNP or ADP solution with fitted linear curves. KLP-6-GFP_AMP-PNP(w) and KLP-6-GFP_AMP-PNP(s) represent weak and strong binding behaviors of KLP-6-GFP in the AMP-PNP solution, respectively. Each plot represents mean ± SEM. In the AMP-PNP solution, 62-82 particles were analyzed, except for KLP-6-GFP_AMP-PNP(w), for which 29 particles were analyzed. In the ADP solution, 115-274 particles were analyzed.

In the presence of ADP, KLP-6-GFP exhibited high diffusion (Fig. 2B and C), but the diffusion coefficient *D*_ADP_ was half of that in the ATP solution (Table 1). Extraneous ADP might not facilitate the complete autoinhibition of KLP-6-GFP as effectively as ADP generated from hydrolysis. KLP-6(1-390)-GFP and KLP-6(D458A)-GFP exhibited higher diffusion than in the ATP solution (Fig. 2B, C, and Table 1). *D*_ADP_ of KLP-6(D458A)-GFP was lower than KLP-6(1-390)-GFP, indicating that the tail domain suppresses the diffusion of KLP-6 in the weak binding state when autoinhibition is relieved.

### Activated full-length KLP-6 is a monomer in solution

Size exclusion chromatography and mass photometry have shown that KLP-6-GFP is a monomeric protein in solution (12). Most kinesins exhibit robust processive movement along microtubules when they form dimers (12, 33). As KLP-6(D458A)-GFP showed processive movement on microtubules, we next analyzed the oligomeric state of this motor using the same methods. Size exclusion chromatography revealed that the peak fraction of KLP-6(D458A)-GFP was equivalent to that of monomeric KLP-6-GFP (Fig. 3A, S1A and B). Mass photometry analyzed this peak fraction, and the molecular weight (125 ± 19 kDa) was consistent with the estimated monomer size (134 kDa) (Fig. 3B). We have previously shown that KLP-6(1-587) and KLP-6(1-587)(D458A) are monomers in solution (12). Given these results, the MBS and MATH domains do not facilitate dimerization when autoinhibition is disrupted. Processive KLP-6(D458A)-GFP is incapable of forming dimers in solution, similar to autoinhibited wild-type KLP-6.

**Figure 3:**
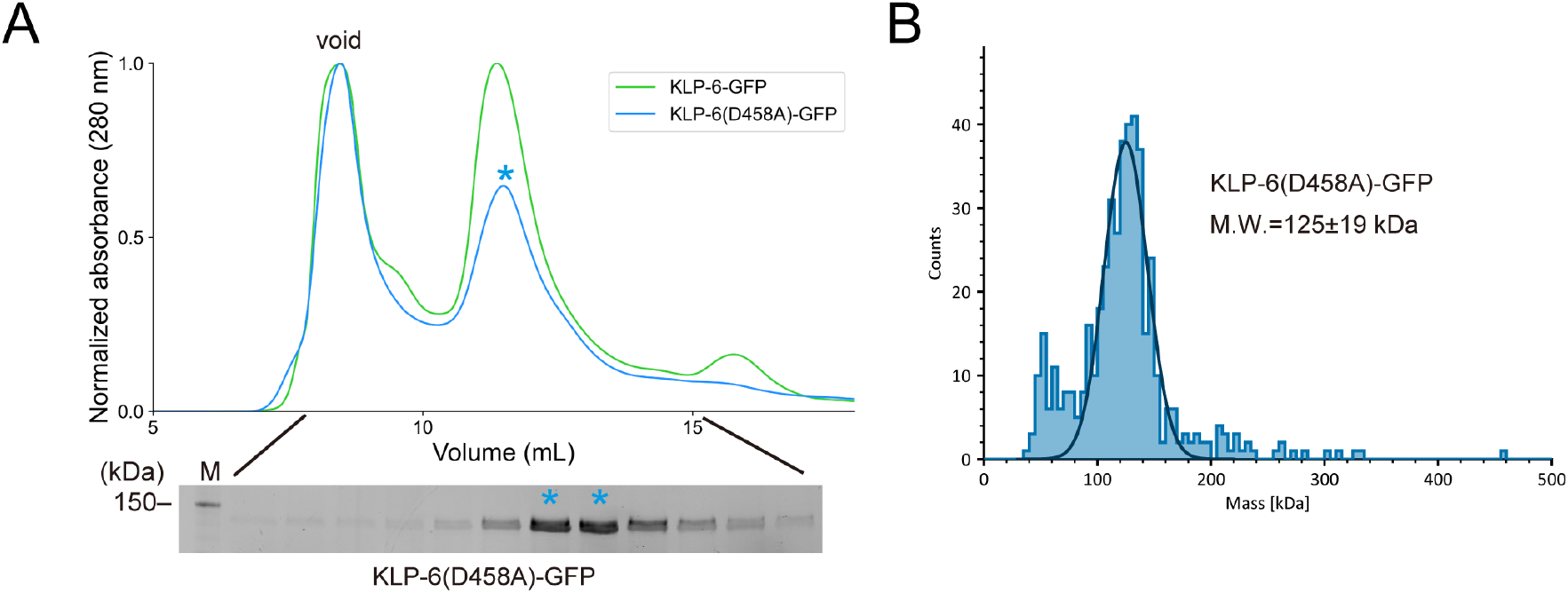
Oligomeric state of KLP-6(D458A)-GFP in solution. (A) Size exclusion chromatography of KLP-6(D458A)-GFP. As a control, the result of KLP-6-GFP from our previous study (12) is shown. The SDS-PAGE of the elution fractions is shown beneath the profile. Asterisks indicate fractions used for mass photometry and single-molecule assays. The void volume of the column is indicated. The number shown on the left side indicates a molecular weight standard. (B) Mass photometry of KLP-6(D458A)-GFP. Histogram shows the particle count of KLP-6(D458A)-GFP at 20 nM. The line represents a Gaussian fit (mean ± SD).

### Activated full-length KLP-6 is a monomer even on microtubules

While some kinesin-3 motors are monomeric in solution, they can dimerize and move processively on microtubules, likely due to the increased local concentration of motors on the microtubule surface (12, 25). We next analyzed the oligomeric states of KLP-6(D458A)-GFP on microtubules using two methods. First, the fluorescence intensities of sfGFP attached to KLP-6 proteins were measured with TIRF microscopy. As a control, the fluorescence intensities of KIF1A(1-393)LZ-GFP, artificially dimerized via the leucine zipper domain, were analyzed. The distribution of fluorescence intensities showed two peaks, likely corresponding to the intensities of one and two sfGFP molecules, respectively (Fig. 4A). It has been reported that some GFP molecules are inactive in solution (34), leading to a partial underestimation of the number of GFP molecules attached to a protein of interest in this assay (35). In contrast to KIF1A(1-393)LZ-GFP, KLP-6-GFP, KLP-6(1-390)-GFP, and KLP-6FL(D458A)-GFP showed just a single peak (Fig. 4A). The fluorescence intensity analysis indicates that these three motors are monomers on microtubules.

**Figure 4:**
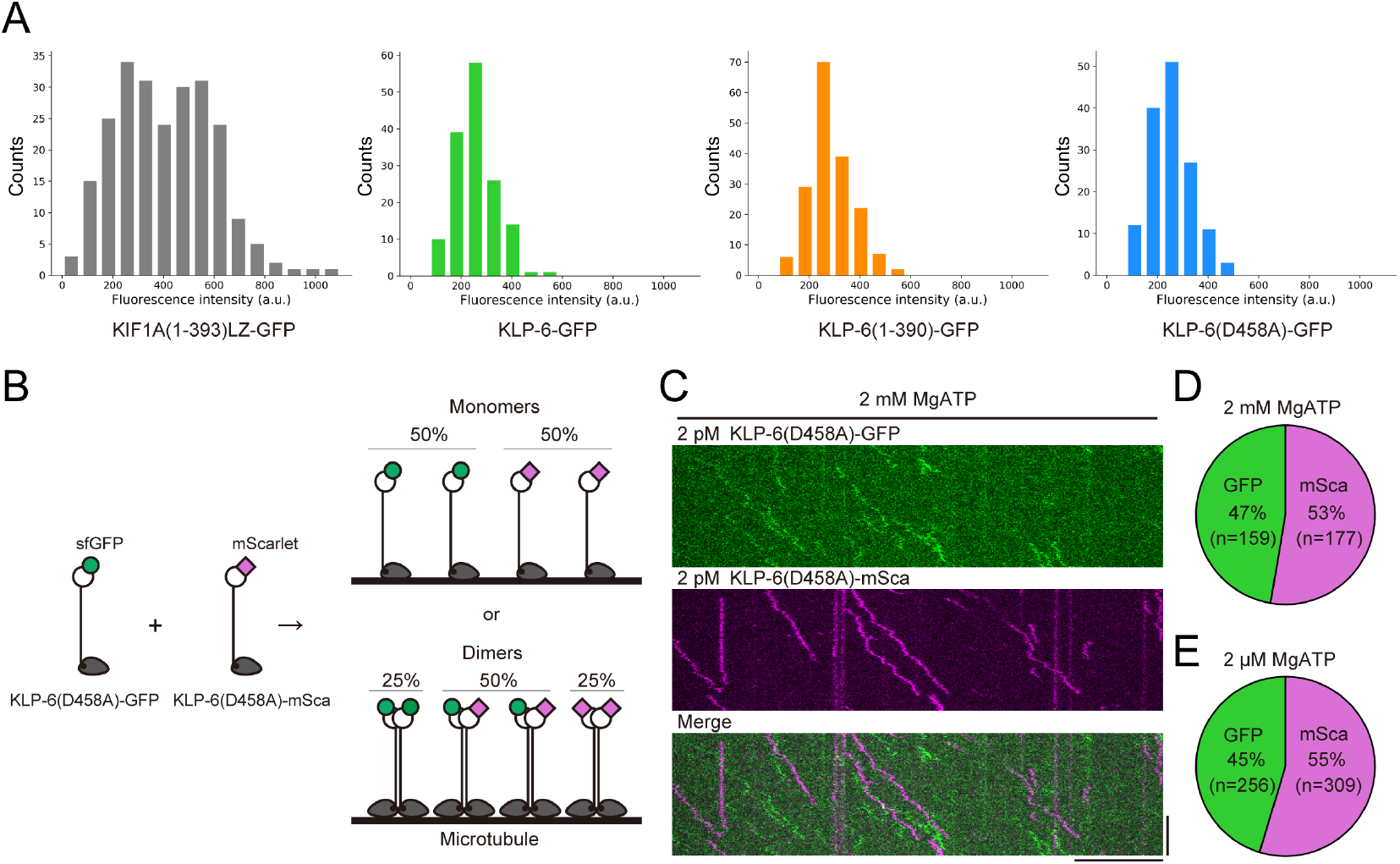
Oligomeric states of KLP-6 proteins on microtubules. (A) Fluorescence intensity distributions of KIF1A(1-393)LZ-GFP, KLP-6-GFP, KLP-6(1-390)-GFP, and KLP-6(D458A)-GFP on microtubules. 144-236 particles were analyzed. (B) Schematic drawing of the dual-color single-molecule assay. If KLP-6(D458A) forms a dimer, 50% co-movement of KLP-6(D458A)-GFP and KLP-6(D458A)-mSca should be observed. (C) Representative kymographs showing the motility of KLP-6(D458A)-GFP and KLP-6(D458A)-mSca observed simultaneously in the presence of 2 mM MgATP. Horizontal and vertical bars represent 10 *μ*m and 60 s, respectively. (D and E) Pie charts displaying the relative populations of KLP-6(D458A)-GFP and KLP-6(D458A)-mSca movements in the presence of 2 mM ATP (D) and 2 *μ*M ATP (E). No co-movements of KLP-6(D458A)-GFP and KLP-6(D458A)-mSca were observed.

Second, full-length KLP-6(D458A) fused with mScarlet (KLP-6(D458A)-mSca) was purified (Figs. S1A and B) and simultaneously observed with KLP-6(D458A)-GFP. If KLP-6(D458A) is able to form dimers, colocalization of GFP and mScarlet caused by heterodimerization between KLP-6(D458A)-GFP and KLP-6(D458A)-mSca should be observed (Fig. 4B). However, such colocalization was not observed, and each fluorescent protein moved independently along microtubules (Fig. 4C). KLP-6(D458A)-GFP and KLP-6(D458A)-mSca had the similar landing ability on microtubules (Fig. S1D), and they were observed in equal proportions in the mixed condition (Fig. 4D and E). These results support the conclusion that KLP-6(D458A) can move processively along microtubules in a monomeric state.

Oligomeric states were determined by size exclusion chromatography, mass photometry, and single-molecule assay. Size exclusion chromatography and mass photometry for KLP-6-GFP and KLP-6(1-390)-GFP were performed in our previous study (12). For KLP-6(580-928)-GFP, only size exclusion chromatography was conducted (Fig. S1A and C). Instantaneous velocities were directly calculated from the raw data and are presented as mean ± SEM; values in parentheses are *p* values for the probability of being wrong in concluding that the mean is not zero (null hypothesis). The minus symbol refers to the polarity of microtubules. Diffusion coefficients *D*_ATP_ and drift velocities were determined from MSD analysis in the presence of 2 mM MgATP. Diffusion coefficients *D*_ADP_ and *D*_AMP−PNP_ were determined from MSD analysis in the presence of 5 mM MgADP and 2 mM MgAMP-PNP, respectively. For KLP-6-GFP, the diffusion coefficients of strong and weak binding states in the AMP-PNP solution are denoted by (s) and (w). All data from MSD analysis are presented as mean ± SEM.

### A Brownian ratchet model explains the processivity of the monomeric KLP-6

The prevailing model for the unidirectional motion of monomeric kinesins along microtubules is described by asymmetric, periodic potentials (10, 14, 15). However, this model does not apply to KLP-6(D458A)-GFP because its motor domain does not have an asymmetric potential, as KLP-6(1-390)-GFP does not show biased movements (Fig. 1B-D). Here, we propose a Brownian ratchet model that considers symmetric potentials and conformational changes in the length between the head and tail domains caused by neck-linker docking (Fig. 5). In this model, the motor and tail domains are approximated as Brownian particles connected by a rigid rod. We assume that the tail domain always follows the motor domain and never overtakes it. The motor domain undergoes conformational changes in response to different nucleotide states, causing the potential to switch between “on” (when the motor domain is nucleotide-free or bound to ATP) and “off” (when the motor domain is bound to ADP) (36, 37). When the potential is on, the motor domain is strongly bound to the microtubule and trapped near one of the potential minima (states S_u_ and S_d_, left panels in Fig. 5). This potential barrier is sufficiently high that the motor cannot overcome it due to thermal noise. When the potential is off, the motor domain is weakly bound to the microtubule, allowing the complex particles to diffuse freely due to thermal noise (states W_d_ and W_u_, right panels in Fig. 5). The tail domain is not directly affected by this potential but its movement is regulated by the rod connected to the motor domain.

**Figure 5:**
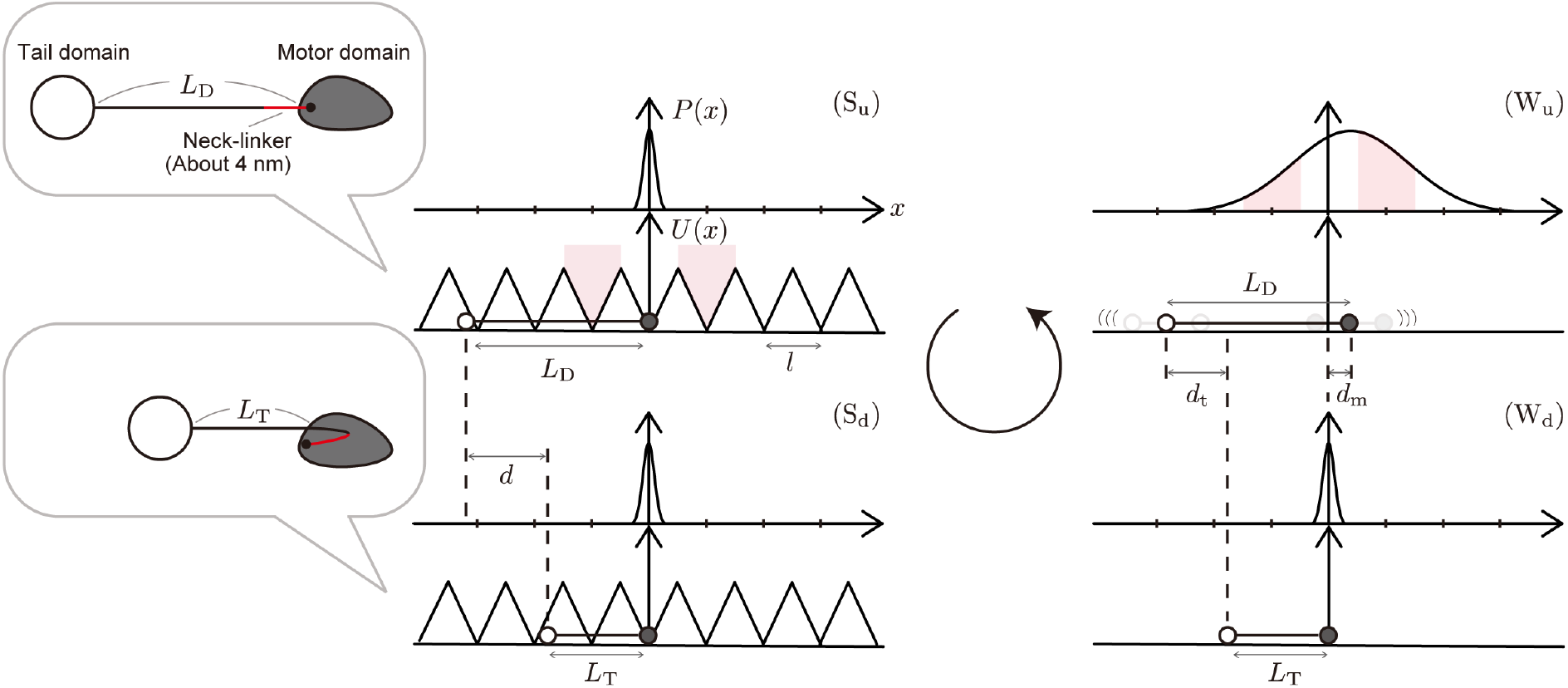
A Brownian ratchet model for activated monomeric KLP-6. The motor domain and tail domain are approximated as Brownian particles connected by a rigid rod. The potential *U*(*x*) for the interaction between the motor domain and microtubule and the probability distribution of the motor domain *P*(*x*) are depicted. When the symmetric, periodic potential of period *l* is on, the motor domain is strongly bound to the microtubule, and trapped near one of the potential minima (left panels, states S_u_ and S_d_). When the potential is off, the motor domain is weakly bound to the microtubule and the complex particles diffuse freely (right panels, states W_u_ and W_d_). In state S_u_ (upper left panel), which is realized right after the potential is switched on, the neck-linker is undocked from the motor domain and the rod length is *L*_D_. This state is short-lived and quickly terminated by the neck-linker docking (transition S_u_ → S_d_), which reduces the rod length to *L*_T_ by *d*. After a while, the potential is switched off by hydrolysis (S_d_ → W_d_), and the release of the neck-linker from the motor domain quickly follows (W_d_ → W_u_). The rod length returns to *L*_D_, causing the motor and tail domains to move forward by *d*_m_ and backward by *d*_t_, respectively. In state W_u_, the complex particles diffuse freely, and the probability distribution broadens symmetrically about the position displaced forward by *d*_m_ from the potential minimum where the motor domain was trapped in the previous strong-binding state. This displacement makes it more likely for the motor domain to be trapped in the forward potential minimum (forward red zone) than in the backward one (backward red zone) in the next strong-binding state.

This model works as follows (the derivation of the expressions given below is provided in Section S3 of the supporting materials and methods). We start with state S_u_ (upper left panel in Fig. 5), which is short-lived and realized immediately after the potential is switched on. In this state, the neck-linker is undocked and the rod length is *L*_D_. The transition S_u_ → S_d_, associated with the neck-linker docking in the motor domain, occurs quickly, reducing the rod length to *L*_T_ and causing the tail domain to move closer to the stationary motor domain by *d*. Right after hydrolysis, the potential is switched off (S_d_ → W_d_), and the release of the neck-linker from the motor domain quickly follows (W_d_ → W_u_). As a result, the rod length returns to *L*_D_, making the motor and tail domains move forward by *d*_m_ and backward by *d*_t_, respectively, according to the law of action and reaction. *d*_m_ and *d*_t_ are given as:

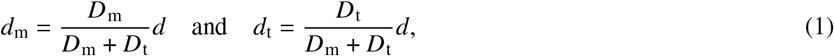

where *D*_m_ and *D*_t_ are the diffusion coefficients of the motor and tail domains in the weak-binding state, respectively. The diffusion coefficient *D*_ADP_ of the complex particles in state W_u_ is given by

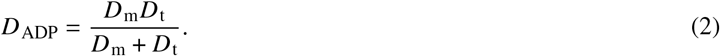

Due to diffusion, the probability distribution function for the motor domain broadens symmetrically about the position displaced forward by *d*_m_ from the potential minimum where the motor domain was trapped in the previous strong-binding state. This displacement makes it more likely for the motor domain to be trapped in a forward potential minimum rather than a backward one in the next strong-binding state. Once the motor domain is trapped in a new or the original potential minimum, another cycle of transitions begins. The drift velocity *v* and diffusion coefficient *D*_ATP_ of the motor undergoing this cycle are expressed as follows:

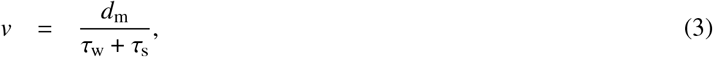

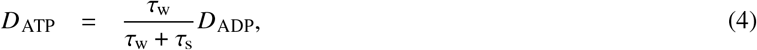

where *τ*_w_ and *τ*_s_ are the dwell times of the strong and weak binding states, respectively. The expression (Eq. 4) is consistent with the diffusion coefficient of the general biased Brownian ratchet motor (10, 14, 38, 39).

Now, we analyze the data of KLP-6(D458A)-GFP listed in Table 1 using Eqs. 2, 3, and 4. Substituting the diffusion coefficient of KLP-6(1-390)-GFP in the ADP solution (*D*_m_ = 0.038 ± 0.008 *μ*m^2^/s) and that of KLP-6(580-928)-GFP (*D*_t_ = 0.189 ± 0.010 *μ*m^2^/s) into Eq. 2, we calculate *D*_ADP_ = 0.032 ± 0.006 *μ*m^2^/s, which is nearly consistent with the observed value of KLP-6(D458A)-GFP (0.025 ± 0.001 *μ*m^2^/s). Next, we validate Eq. 3 for drift velocity. The neck-linker length is almost 4 nm (26, 40), and when it fully binds to the motor domain, *d* can be up to 8 nm, as shown in Fig. 5. If *d* = 6 or 8 nm, the hydrolysis rate *k* = 1/(*τ*_w_ + *τ*_s_) = 66.9 /s or 50.2 /s, respectively, is required for KLP-6(D458A)-GFP to move directionally on microtubules at 67.2 nm/s. This hydrolysis rate is consistent with a previous study (26) which showed that KLP-6(D458A) has a similar hydrolysis rate to the well-known kinesin-1 motor domain (about 60 /s) (10, 17, 41– 43). According to Eq. 4, the ratio *D*_ADP_/*D*_ATP_ is governed by the dwell time of each state and is not affected by biased movement, similar to the well-known biased Brownian ratchet motor. (10, 13, 14, 16, 38, 39). We analyzed KLP-6(1-390)-GFP and KLP-6(D458A)-GFP and found that the ratio *D*_ADP_/*D*_ATP_ of KLP-6(D458A)-GFP, which is 2.78 ± 1.24, is 2.5 times larger than that of KLP-6(1-390)-GFP, which is 1.12 ± 0.24. This result indicates that either *τ*_w_ or *τ*_s_, or both, of KLP-6(D458A)-GFP are significantly changed from those of KLP-6(1-390)-GFP due to the addition of the stalk and tail domains, which will be discussed later.

## DISCUSSION

### Autoinbition and diffusion of KLP-6

Our MSD analysis provides valuable insights into the autoinhibition and diffusion of KLP-6. In the AMP-PNP solution, we observed the transition of KLP-6-GFP from a weak-binding state to a strong-binding state (Fig. 2A), likely corresponding to the shift from the autoinhibited state to the activated state. This result indicates that the combination of AMP-PNP (or ATP) and microtubules helps autoinhibited KLP-6-GFP release ADP from the motor domain, thereby alleviating autoinhibition (26). However, the diffusion coefficient of KLP-6-GFP is similar to that of KLP-6(580-928)-GFP in the presence of ATP (Table 1), suggesting it quickly returns to the highly diffusive autoinhibited state as soon as ADP is generated after hydrolysis. The diffusion coefficients *D*_ATP_ and *D*_ADP_ of KLP-6(D458A)-GFP are much smaller than those of KLP-6-GFP and KLP-6(580-928)-GFP (Table 1), indicating that the D458A mutation almost completely disrupts the autoinhibition of KLP-6. Our model supports this finding, as *D*_ADP_ of KLP-6(D458A)-GFP can be explained without considering the highly diffusive autoinhibited state.

The ratio *D*_ADP_/*D*_ATP_ of KLP-6(D458A)-GFP is significantly larger than that of KLP-6(1-390). This ratio depends on the dwell time of each nucleotide state, as shown in Eq. 4. One potential explanation is that the stalk domain acts as a steric hindrance to ATP binding to the motor domain, prolonging the apo state. Another possibility is that the tail domain suppresses the diffusion of KLP-6(D458A)-GFP in the ADP state, making it easier for the motor domain to find suitable positions on microtubules to release ADP. Indeed, in the case of KIF1A, the L7 loop interacts with *β*-tubulin, facilitating the release of ADP (44).

### Application of our model to other motors

Our model differs significantly from the well-known Brownian ratchet model (10, 13–16, 39) and suggests that neck-linker docking and the second microtubule-binding domain, rather than the asymmetric potential, are essential for the movement of KLP-6(D458A)-GFP. Similar to KLP-6(D458A)-GFP, the movement of K351-BDTC might be driven by neck-linker docking in addition to the asymmetric potential (17). Our model estimated that if *d* = 8 nm and *D*_ADP_ of K351 is twice as large as that of BDTC, K351-BDTC can move four times faster than K351, as observed experimentally (17) (see Section S4 of the supporting materials and methods and Fig. S2). Indeed, BDTC is likely less diffusive than K351 because the diffusion coefficient of K351-BDTC is half that of the two-headed kinesin fragment, K411 (17).

Our model may not be applicable to Kar3 with a non-catalytic head. Kar3 remains slightly diffusive even in the ATP and apo states (19), but heterodimerization with the non-catalytic Cik-1 or Vik-1 enables Kar3 to bind strongly to microtubules, leading to an asymmetric potential (45, 46). ATP binding at Kar3 drives a large rotation of the coiled-coil stalk, causing the non-catalytic domain to detach from the microtubule (47, 48). When the non-catalytic domain is dissociated, the tension from the stalk domain attempting to return to its original length, as proposed by our model, does not cause Kar3 to move forward along the microtubule.

For heterodimers composed of processive and non-processive kinesins, as shown in previous studies (20–22), it is important to consider the applicability of our model. A MINFLUX study of kinesin has shown that conventional kinesin takes both chassé-inchworm steps and hand-over-hand steps (49). During chassé-inchworm steps, the front processive head may be accelerated by tension from the stalk domain, given that the front processive head and rear non-processive head correspond to the roles of the motor domain and tail domain of KLP-6(D458A), respectively. Conversely, if they take hand-over-hand steps, such tension would not accelerate the front head because the non-processive head overtakes the front head.

It is uncertain whether KLP-6 functions as a monomer in cells, but dimerization is expected to be necessary for efficient cargo transport. This is because the speed of monomeric KLP-6 is about four-fold and ten-fold slower than that of artificially dimerized KLP-6(1-390) in vitro (0.26±0.12 *μ*m/s) (12) and GFP-labeled KLP-6 in cilia (0.72±0.18 *μ*m/s) (28), respectively. Nevertheless, we believe the analysis of the KLP-6 monomer presented in this study provides a good model for understanding the processivity of the kinesin motor domain. Furthermore, it will be helpful for creating rational artificial molecular motors since many of them rely on thermal fluctuations to generate unidirectional movements. (50). Our model concludes that not only the asymmetric potential but also slight conformational change gives monomeric motors robust unidirectional movement.

## SUPPORTING MATERIAL

Supporting material can be found online.

## AUTHOR CONTRIBUTIONS

T.K., K.S. and S.N. designed research; T.K and S.N. performed experiments; K.S. performed calculations; T.K analyzed data; T.K., K.S. and S.N. wrote the paper.

## ACKNOWLEDGMENTS

Generous support from the FRIS CoRE, which is a shared research environment. We also would like to thank Dr. Atsushi Nakagawa and Mr. Jiye Wang (Osaka University) for technical assistance. TK was supported by JSPS KAKENHI (grant no. JP23KJ0168). SN was supported by JSPS KAKENHI (grant no. JP23H02472). This work was performed under the Collaborative Research Program of Institute for Protein Research, Osaka University, CR-24-02.

## STATEMENT

During the preparation of this work, the authors used GPT-4 to check English grammar and enhance their writing. After using this tool, the authors reviewed and edited the content as necessary and take full responsibility for the final publication.

## SUPPORTING CITATIONS

References (51–53) are cited in the supporting material.

