## Supplemental information for "Processivity of the monomeric KLP-6 kinesin and a Brownian ratchet model with symmetric potentials"

#### SUPPORTING MATERIALS AND METHODS

#### S1 Expression and purification of recombinant proteins

Sf9 cells (Thermo Fisher Scientific) were maintained in Sf900 II SFM (Thermo Fisher Scientific) at 27°C. DH10Bac (Thermo Fisher Scientific) was transformed to generate bacmid. To prepare baculovirus, $1 \times 10^6$ cells of Sf9 cells were transferred to each well of a tissue culture-treated six-well plate. After the cells attached to the bottom of the dishes, about ~ 5 $\mu$g of bacmid were transfected using 5 $\mu$L of TransIT-Insect transfection reagent (Takara Bio Inc). Five days after initial transfection, the culture media were collected and spun at 3000 × g for 3 min to obtain the supernatant (P1). For protein expression, 400 mL of Sf9 cells ($2 \times 10^6$ cells/mL) were infected with 200 $\mu$L of P1 virus and cultured for 65 hr at 27°C. Sf9 cells were resuspended in 25 mL of protein buffer (50 mM HEPES-KOH, pH 7.5, 150 mM KCH3COO, 2 mM MgSO4, 1 mM EGTA, 10% glycerol) along with 1 mM DTT, 1 mM PMSF, 0.1 mM ATP, and 0.5% Triton X-100. After incubating on ice for 10 min, lysates were cleared by centrifugation (15,000 × g, 20 min, 4°C) and loaded on Streptactin-XT resin (IBA Lifesciences, Göttingen, Germany) (bead volume: 2 mL). The resin was washed with 40 mL Strep wash buffer (50 mM HEPES-KOH, pH 8.0, 450 mM $KCH_3COO$, 2 mM MgSO4, 1 mM EGTA, 10% glycerol). Protein was eluted with 40 mL Strep elution buffer (50 mM HEPES-KOH, pH 8.0, 150 mM KCH3COO, 2 mM MgSO4, 1 mM EGTA, 10% glycerol, 300 mM biotin). Eluted solution was concentrated using an Amicon Ultra 15 (Merck) and then separated on an NGC chromatography system (Bio-Rad) equipped with a Superdex 200 Increase 10/300 GL column (Cytiva). Peak fractions were collected and concentrated using an Amicon Ultra 4 (Merck). Concentrated proteins were aliquoted and snap-frozen in liquid nitrogen. KIF5A(1-416)::mScarlet::Strep-tag II and KIF1A(1-393)LZ::sfGFP::His-tag were expressed in bacteria and purified as previously described (1).

#### S2 Total internal reflection fluorescence single-molecule motility assays

Tubulin was purified from porcine brain as described (2). Tubulin was labeled with Biotin-PEG2-NHS ester (Tokyo Chemical Industry, Tokyo, Japan) and AZDye647 NHS ester (Fluoroprobes, Scottsdale, AZ, USA) as described (3). To polymerize Taxol-stabilized microtubules labeled with biotin and AZDye647, 30 $\mu$M unlabeled tubulin, 1.5 $\mu$M biotin-labeled tubulin, and 1.5 $\mu$M AZDye647-labeled tubulin were mixed in BRB80 buffer supplemented with 1 mM GTP and incubated for 15 min at 37°C. Then, an equal amount of BRB80 supplemented with 40 $\mu$M taxol was added and further incubated for more than 15 min. The solution was loaded on BRB80 supplemented with 300 mM sucrose and 20 $\mu$M taxol and ultracentrifuged at 100,000 × g for 5 min at 30°C. The pellet was resuspended in BRB80 supplemented with 20 $\mu$M taxol. TIRF assays using the purified porcine microtubules were performed as described (4). Glass chambers were prepared by acid washing as previously described (5). Glass chambers were coated with PLL-PEG-biotin (SuSoS, Dübendorf, Switzerland). Polymerized microtubules were flowed into streptavidin adsorbed flow chambers and allowed to adhere for 5–10 min. Unbound microtubules were washed away using assay buffer (90 mM HEPES-KOH pH 7.4, 50 mM KCH3COO, 2 mM Mg(CH3COO)2, 1 mM EGTA, 10% glycerol, 0.1 mg/mL biotin-BSA, 0.2 mg/ mL kappa-casein, 0.5% Pluronic F127, nucleotide of interest (ATP, ADP, or AMP-PNP) diluted to indicated concentrations, and an oxygen scavenging system composed of PCA/PCD/Trolox). Purified motor proteins were diluted to concentrations suitable for observing single particles (2-150 pM) in the assay buffer. Diluted KIF5A(1-416)-mScarlet was added to the assay buffer to determine the polarity of the microtubules. The solution was then introduced into the glass chamber. An ECLIPSE Ti2-E microscope equipped with a CFI Apochromat TIRF 100XC Oil objective lens (1.49 NA), an Andor iXion life 897 camera and a Ti2-LAPP illumination system (Nikon, Tokyo, Japan), was used to observe single molecule motility. NIS-Elements AR software version 5.2 (Nikon) was used to control the system.

#### S3 A model for monomeric KLP-6

We consider the Brownian ratchet model shown in Fig. 5 of the main text for activated full-length KLP-6. The motor and tail domains are approximated as Brownian particles connected by a rigid rod. When the potential is off (states $W_d$ and $W_u$), the dynamics of the motor domain (m) and tail domain (t) are described by the following overdamped Langevin equations:

$$\gamma_m \frac{dx_m}{dt} = f(t) + \xi_m(t), \tag{S1}$$

$$\gamma_t \frac{dx_t}{dt} = -f(t) + \xi_t(t), \tag{S2}$$

where $\gamma_i$ ($i = \mathrm{m}, \mathrm{t}$) is the coefficient of viscous drag, $f(t)$ is the force on the motor domain from the tail domain via the rod, and $\xi_i$ is the Gaussian white noise which satisfies the following fluctuation dissipation theorem:

$$\langle \xi_i(t) \rangle = 0, \quad \langle \xi_i(t) \xi_j(t') \rangle = 2\gamma_i k_\mathrm{B} T \delta_{ij} \delta(t - t') \tag{S3}$$

where $\langle \cdots \rangle$ indicates the statistical average, $k_\mathrm{B}$ is the Boltzmann constant, and $T$ is the temperature. The diffusion coefficient $D_i$ satisfies $D_i = k_\mathrm{B}T/\gamma_i$. The Langevin equation for "the center of viscosity" $X$, which is a counterpart of the center of mass in the Newtonian mechanics, of the two particles defined by

$$X = \frac{\gamma_\mathrm{m} x_\mathrm{m} + \gamma_\mathrm{t} x_\mathrm{t}}{\gamma_\mathrm{m} + \gamma_\mathrm{t}} = \frac{D_\mathrm{t} x_\mathrm{m} + D_\mathrm{m} x_\mathrm{t}}{D_\mathrm{t} + D_\mathrm{m}}, \tag{S4}$$

is obtained from Eqs. S1 and S2 as follows:

$$\gamma_X \frac{dX}{dt} = \xi_X(t), \tag{S5}$$

where

$$\gamma_X = \gamma_\mathrm{m} + \gamma_\mathrm{t}, \quad \xi_X = \xi_\mathrm{m} + \xi_\mathrm{t}, \tag{S6}$$

and $\xi_X$ satisfies Eq. S3 with $i = X$. From Eq. S5, we see that the diffusion coefficient for the center of viscosity is given by

$$D_X = \frac{k_\mathrm{B} T}{\gamma_X} = \frac{D_\mathrm{m} D_\mathrm{t}}{D_\mathrm{m} + D_\mathrm{t}}, \tag{S7}$$

which corresponds to Eq. 2 in the main text.

Suppose that the motor is in state $\mathrm{S_d}$ (lower left panel of Figure 5 in the main text) and the motor domain is trapped near the potential minimum at $x = 0$. At time $t = 0$, the potential is switched off (transition $\mathrm{S_d} \longrightarrow \mathrm{W_d}$). In this situation, the center of viscosity is localized near $x = X_0$ at $t = 0$ where

$$X_0 = -\frac{D_\mathrm{m}}{D_\mathrm{m} + D_\mathrm{t}} L_\mathrm{T}, \tag{S8}$$

since $x_\mathrm{m} \approx 0$ and $x_\mathrm{t} = x_\mathrm{m} - L_\mathrm{T}$ at this moment. Therefore, the probability density function $P_\mathrm{cv}(X, t)$ for the center of viscosity $X$ at $t > 0$ is given by

$$P_\mathrm{cv}(X, t) = P_0(X - X_0, t), \tag{S9}$$

where $P_0(x, t)$ is the normal distribution of zero mean and variance $2D_X t$:

$$P_0(x, t) = \frac{1}{2\sqrt{\pi D_X t}} \exp\left( - \frac{x^2}{4 D_X t} \right). \tag{S10}$$

We assume that the rod length increases from $L_\mathrm{T}$ to $L_\mathrm{D}$ by $d$ ($\mathrm{W_d} \longrightarrow \mathrm{W_u}$) right after the potential is switched off ($2D_\mathrm{X} t \ll l^2$ with $l$ being the period of the potential) before the center of viscosity does not diffuse appreciably. It is clear from the definition of $X$ given in Eq. S4 that $x_\mathrm{m} - X$ and $X - x_\mathrm{t}$ increase by

$$d_\mathrm{m} \; = \; \frac{D_\mathrm{m}}{D_\mathrm{m} + D_\mathrm{t}} d \quad \text{and} \quad d_\mathrm{t} \; = \; \frac{D_\mathrm{t}}{D_\mathrm{m} + D_\mathrm{t}} d, \tag{S11}$$

respectively, as a result of this conformational change. This means that the motor and tail domains move forward by $d_\mathrm{m}$ and backward by $d_\mathrm{t}$, respectively, in response to the conformational change, since the center of viscosity remains localized around $x = X_0$ during this process. Eq. S11 corresponds to Eq. 1 in the main text.

In state $\mathrm{W_u}$ (upper right panel of Fig. 5 in the main text), the motor domain is a distance

$$L_\mathrm{m} = \frac{D_\mathrm{m}}{D_\mathrm{m} + D_\mathrm{t}} L_\mathrm{D} \tag{S12}$$

ahead of the center of viscosity $X$. Therefore, the distribution $P(x_\mathrm{m}, t)$ of the motor domain position $x_\mathrm{m}$ in this state is obtained from that of $X$ as

$$P(x_\mathrm{m}, t) = P_\mathrm{cv}(x_\mathrm{m} - L_\mathrm{m}, t) = P_0(x_\mathrm{m} - d_\mathrm{m}, t), \tag{S13}$$

where the last equality follows from Eqs. S8, S9, S11, and S12; and $P_0(x, t)$ is the normal distribution defined in Eq. S10.

After the dwell time of the weak binding state ($\tau_\mathrm{w}$) passes since the potential is switched off, it is switched on again, and the motor domain is trapped in one of the potential valleys ($\mathrm{W_u} \longrightarrow \mathrm{S_u}$). Let $p_n$ be the probability that the motor domain is displaced by $nl$ (where $n$ is an integer and $l$ is the period of the potential) from $x = 0$, where it was previously trapped. The displacement $nl$ is realized if $x_\mathrm{m}$ is in the interval $(nl - l/2, nl + l/2)$ right before the potential is switched on at $t = \tau_\mathrm{w}$. Therefore, we have

$$p_n = \int_{nl-l/2}^{nl+l/2} P(x_\mathrm{m}, \tau_\mathrm{w}) dx_\mathrm{m} = \int_{nl-l/2-d_\mathrm{m}}^{nl+l/2-d_\mathrm{m}} P_0(x, \tau_\mathrm{w}) dx, \tag{S14}$$

where we have used Eq. S13. For the highly diffusive motors, $2D_X\tau_\mathrm{w} \gg l^2$ is satisfied, and the mean $l\langle n\rangle$ and variance $l^2(\langle n^2\rangle - \langle n\rangle^2)$ of the displacement are approximately expressed as (6, 7)

$$l\langle n\rangle = l \sum_{n=-\infty}^{\infty} np_n \approx d_\mathrm{m}, \quad l^2(\langle n^2\rangle - \langle n\rangle^2) \approx 2D_X\tau_\mathrm{w} \tag{S15}$$

Since these are the mean and the variance of the displacement per reaction cycle with duration $\tau_\mathrm{w} + \tau_\mathrm{s}$, where $\tau_s$ is the dwell time of the strong-binding state, the drift velocity $v$ and the diffusion coefficient $D$ of the motor are given by

$$v = \frac{l\langle n\rangle}{\tau_\mathrm{w} + \tau_\mathrm{s}} \approx \frac{d_\mathrm{m}}{\tau_\mathrm{w} + \tau_\mathrm{s}} \tag{S16}$$

and

$$D = \frac{l^2(\langle n^2\rangle - \langle n\rangle^2)}{2(\tau_\mathrm{w} + \tau_\mathrm{s})} \approx \frac{\tau_\mathrm{w}}{\tau_\mathrm{w} + \tau_\mathrm{s}} D_\mathrm{X}, \tag{S17}$$

respectively. Eqs. S16 and S17 correspond to Eqs. 3 and 4 in the main text, respectively.

#### S4 Models for K351 and K351-BDTC

Inoue *et al*. have shown that attaching a non-catalytic microtubule-binding protein, BDTC, to the C-terminal region of monomeric K351 accelerated the motor speed by four times (8). Our model can explain this result. While the processivity of monomeric K351 is explained by the prevailing Brownian ratchet model, which takes into account asymmetric, periodic potentials (Fig. S2A), the processivity of K351-BDTC is explained by a model that considers both asymmetric potentials and a length change between the head and BDTC (Fig. S2B). In this model, K351 and BDTC correspond to the motor domain and tail domain of KLP-6(D458A), respectively. The difference from the model for KLP-6(D458A) is that after the potential is switched on, K351 is trapped in an asymmetric potential well (transition $\mathrm{W_u} \longrightarrow \mathrm{S_u}$). Immediately after this transition, BDTC approaches K351 by $d$ due to neck-linker docking in K351 ($\mathrm{S_u} \longrightarrow \mathrm{S_d}$). Right after hydrolysis, the potential is switched off ($\mathrm{S_d} \longrightarrow \mathrm{W_d}$), and then K351 moves forward by $d_\mathrm{m}$ ($\mathrm{W_d} \longrightarrow \mathrm{W_u}$), as shown in the model for KLP-6(D458A). Both this displacement and the asymmetry of the potential help K351 to be trapped with higher probability in the forward potential well (forward red zone) than in the backward one (backward red zone) in the next strong-binding state. Let $x = 0$ be the position of the potential minimum where K351 was trapped in the previous strong-binding state, $l$ be the period of the potential, and $x = l/2 - d_\mathrm{p}$ be the location of the closest potential maximum on the right of $x = 0$. Then, the probability that K351 is trapped by the potential minimum at $x = nl$, with $n$ being an integer, in the next strong-binding state is given by

$$p_n = \int_{nl-l/2-d_\mathrm{p}}^{nl+l/2-d_\mathrm{p}} P(x_\mathrm{m}, \tau_\mathrm{w})\, dx_\mathrm{m} = \int_{nl-l/2-d_\mathrm{p}-d_\mathrm{m}}^{nl+l/2-d_\mathrm{p}-d_\mathrm{m}} P_0(x, \tau_\mathrm{w})\, dx, \tag{S18}$$

where the probability distributions $P$ and $P_0$ have been defined in Eqs. S13 and S10, respectively, and $\tau_\mathrm{w}$ is the dwell time of the weak-binding state, as before. Comparing the rightmost expressions in Eqs. S14 and S18, we see that $p_n$ for the model of K351-BDTC is obtained from the one for the model of KLP-6(D458A) by replacing $d_\mathrm{m}$ with $d_\mathrm{p} + d_\mathrm{m}$. Therefore, the drift velocity of K351-BDTC is approximately given as:

$$v = \frac{d_\mathrm{p} + d_\mathrm{m}}{\tau_\mathrm{w} + \tau_\mathrm{s}}, \tag{S19}$$

where $\tau_\mathrm{s}$ is the dwell time of the strong-binding state. It should be clear that if we set $d_\mathrm{m} = 0$, the model for K351-BDTC is mathematically equivalent to the one for K351. Hence, the drift velocity of K351 is approximately given as:

$$v = \frac{d_\mathrm{p}}{\tau_\mathrm{w} + \tau_\mathrm{s}}. \tag{S20}$$

Now, we analyze the data from Inoue et al. (8). The hydrolysis rates $k$ for K351 and $k_\mathrm{B}$ for K351-BDTC are 66 /s and 27 /s, respectively. The drift velocities $v$ for K351 and $v_\mathrm{B}$ for K351-BDTC are 40 nm/s and 161 nm/s, respectively. It should be noted that these hydrolysis rates and drift velocities were measured from motors that included immotile ones (8). We calculated $d_\mathrm{p} = v/k = 0.61$ nm and $d_\mathrm{m} = v_\mathrm{B}/k_\mathrm{B} - d_\mathrm{p} = 5.35$ nm. If $d = 8$ nm, the ratio between the diffusion coefficient of K351 in the weak binding state ($D_\mathrm{m}$) and that of BDTC ($D_\mathrm{t}$) is calculated as $D_\mathrm{t}/D_\mathrm{m} = d/d_\mathrm{m} - 1 = 0.50$ from Eq. S11. Indeed, BDTC is likely less diffusive than K351 because the diffusion coefficient of K351-BDTC is half that of the two-headed kinesin fragment, K411 (8).


### SUPPORTING MOVIE AND FIGURES

**Movie S1. An example movie showing the motility of KLP-6(D458A)-GFP** In the first frame, microtubules were imaged in the 640 channel. In the subsequent frames, KLP-6(D458A)-GFP was imaged in the 488 channel. The images from both channels were merged. The movie is played back at 60 × speed.

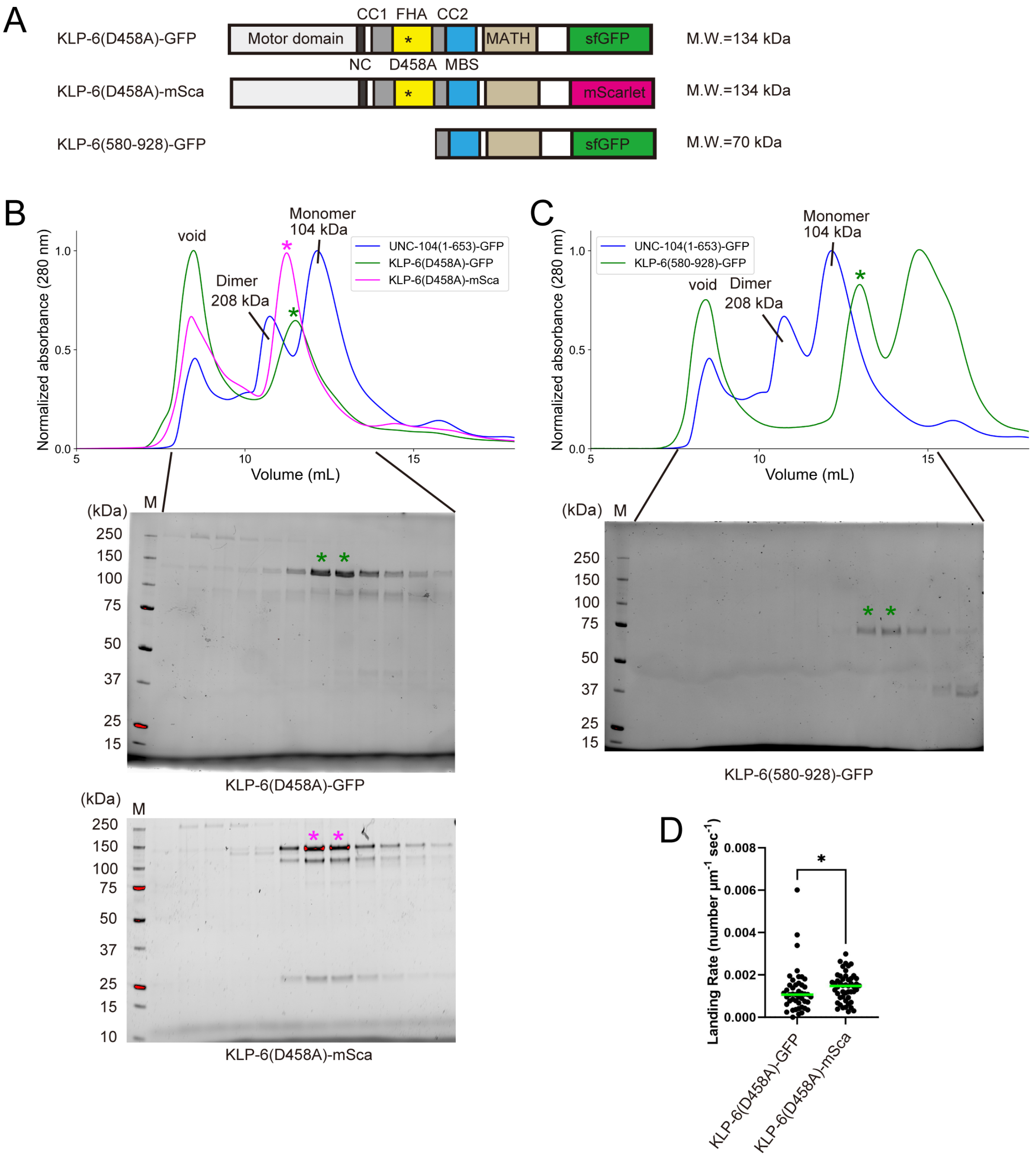


Figure S1: Purification of KLP-6 proteins. (A) Schematic drawing of the domain organization of KLP-6(D458A)-GFP, KLP-6(D458A)-mSca, and KLP-6(580-928)-GFP. Calculated molecular weight is written at the right side. (B) Size exclusion chromatography of KLP-6(D458A)-GFP and KLP-6(D458A)-mSca. As a control, the result of UNC-104(1-653)-GFP reported in our previous study is shown (4). The SDS-PAGE of the elution fractions is shown beneath the profile. Asterisks indicate fractions used for single-molecule assays. The void volume of the column is indicated. (C) Size exclusion chromatography of KLP-6(580-928)-GFP. As a control, the result of UNC-104(1-653)-GFP reported in our previous study is shown (4). The SDS-PAGE of the elution fractions is shown beneath the profile. Asterisks indicate fractions used for single-molecule assays. The void volume of the column is indicated. (D) Dot plots showing the landing rate of 7 pM KLP-6(D458A)-GFP and KLP-6(D458A)-mSca. Each dot shows a single datum point. Green bars represent median value. n=45 and 49 microtubules for KLP-6(D458A)-GFP and KLP-6(D458A)-mSca, respectively. Mann-Whitney U test. *, $P$ <0.05.

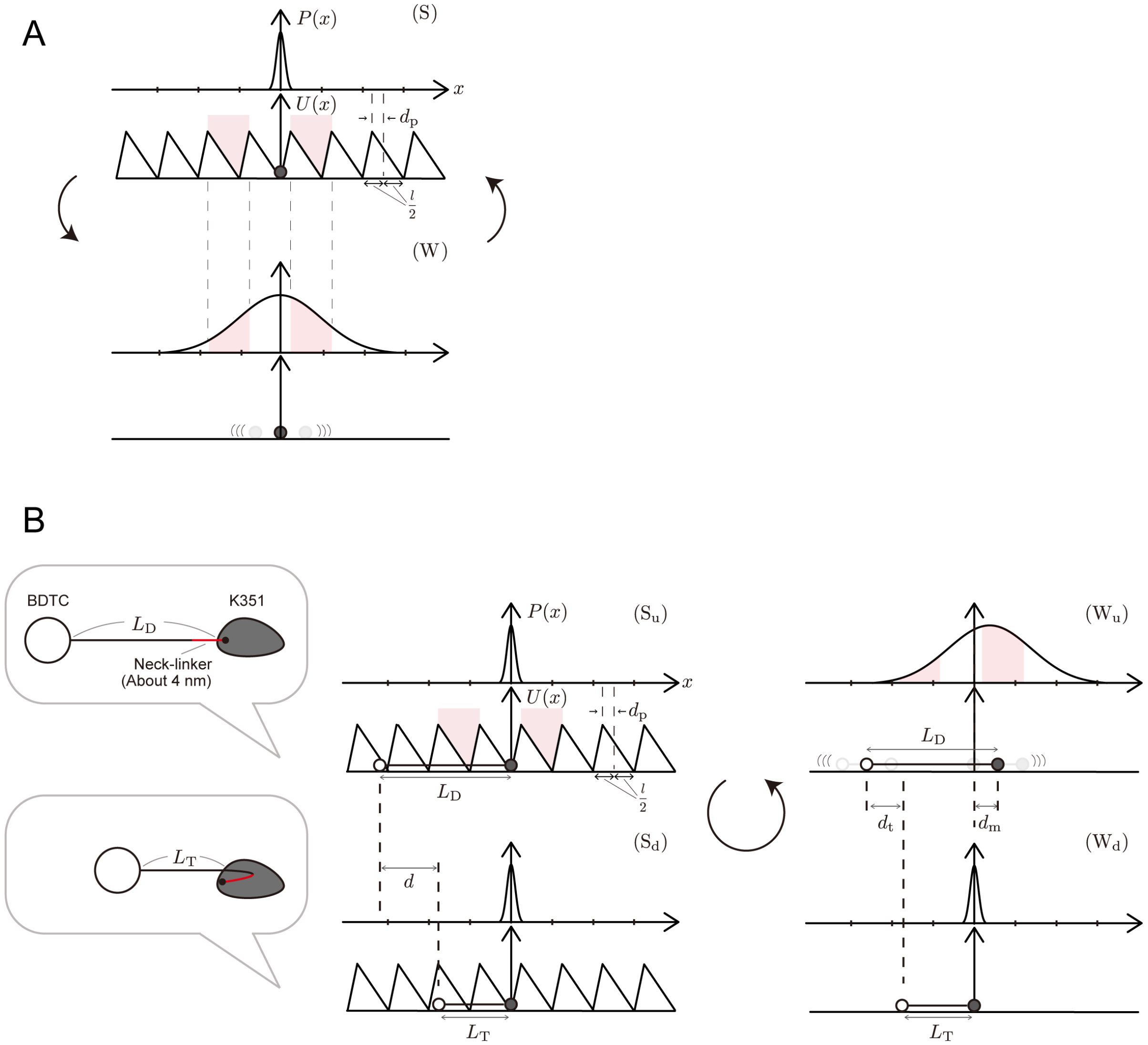


Figure S2: Models for (A) K351 and (B) K351-BDTC. K351 and BDTC are approximated as Brownian particles in the both model, and they are connected by a rigid rod for K351-BDTC in Model B. The asymmetric, periodic potential $U(x)$ for the interaction between K351 and microtubules and the probability distribution $P(x)$ of K351 are depicted. The period of the potential is $l$, and each position of the potential maxima is shifted to the left by $d_p$ from the midpoint between the neighboring minima on the right and left. When the potential is on (state S, upper panel in A; states $S_u$ and $S_d$, left panels in B), K351 is strongly bound to the microtubule and is trapped near one of the potential minima. When the potential is off (state W, lower panel in A; states $W_u$ and $W_d$, right panels in B), K351 is weakly bound to the microtubule, and K351 and K351-BDTC diffuse freely in Models A and B, respectively. Model A: After the potential is switched off (transition S → W), K351 diffuses symmetrically about the potential minimum where it was trapped. Then, when the potential is switched on again, K351 is more likely to be trapped in the forward potential well (forward red zone) than in the backward one (backward red zone) because of the asymmetry of the potential. Model B: When the potential is switched on, the rod length is $L_D$ (state $S_u$) and immediately reduces to $L_T$ due to the neck-linker docking in K351, causing BDTC to approach the stationary K351 by $d$ (state $S_d$). Right after hydrolysis, the potential is switched off ($S_d \to W_d$), and then the rod length returns to $L_D$, causing K351 and BDTC to move forward by $d_m$ and backward by $d_t$, respectively ($W_d \to W_u$). In state $W_u$, K351-BDTC diffuses symmetrically about the point displaced forward by $d_m$ from the potential minimum where it was in the previous strong-binding state. Both this displacement and the asymmetry of the potential help K351 to be trapped with higher probability in the forward potential well (forward red zone) than in the backward one (backward red zone) in the next strong binding state.
